# Rapid population decline of Egyptian fruit bats on Cyprus

**DOI:** 10.64898/2026.06.05.728241

**Authors:** Dina K.N. Dechmann, Karin L. Akre, Luz A. deWit, Artemis G.G. Kafkaletou Diez, Isabella Mandl, Zoe Makridou, Koulla Michael, M. Teague O’Mara, Klitos Papastylianou, Martin Wikelski, Winifred F. Frick

## Abstract

Preventing extirpations requires timely detection of rapid declines and immediate conservation response. Egyptian fruit bats (*Rousettus aegyptiacus*) are globally listed as Least Concern, but their genetically isolated population in Cyprus is Critically Endangered. We document a severe ongoing decline of the Cypriot *R. aegyptiacus* population observed during standardized counts at major roost sites. April 2024 counts at four principal roosts recorded 4633 bats. Surveys in December 2025 documented 554 individuals across the same sites. April 2026 counts revealed a 90.1% decline since April 2024. Additionally, we found numerous dead bats at three roosts, and residents across the island reported dead bats, suggesting an acute mortality event beginning in early 2025. The cause remains unknown. Given the magnitude and pace of decline, this population potentially faces imminent extirpation. We recommend immediate action to protect roosting and foraging habitat, along with coordinated diagnostics, expedited research permits, standardized monitoring, and technical collaboration to identify and mitigate underlying causes of high mortality.

---

Rapid population collapses can lead to irreversible biodiversity loss if not detected and addressed quickly (Woinarski, 2018). This risk is especially high for geographically distinct populations, whose extirpation can reduce genetic diversity and destabilize ecosystems (Ehrlich & Daily, 1993). Early documentation of declines increases the effectiveness of interventions by enabling the mobilization of conservation measures.

We document evidence of a rapid and severe decline of Egyptian fruit bats (*Rousettus aegyptiacus*) on the island of Cyprus through a series of count surveys at key roosting sites to establish recent trends across the island. This species is globally listed as Least Concern (Korine, 2016), but the genetically distinct Cypriot population is Critically Endangered (Russo & Cistrone, 2023) due to historical persecution and its restricted insular distribution. The reporting period for the most recent Article 17 Habitats Directive (Dir. 92/43/EEC) species report ended in 2024, before the mortality event documented here.

Egyptian fruit bats depend on high-humidity caves, or similar structures. In Cyprus, most of the population has recently roosted in 15 shelters across the island (Figure 1; Supplementary Table 1). We conducted standardized exit surveys at known roost sites by counting bats as they emerged at sunset. April 2024 and December 2025 counts were conducted at four caves (exit counts and one internal visual estimate: Mammari, April 2024), with two additional caves in 2025; 2026 counts continued at five shelters, and five shelters were added (Table 1). Exit counts began at sunset and continued until no bats were seen emerging (average duration: 150 min, range 62-270 min). Exit counts were conducted visually in situ, using red light for low light levels, or by video analysis of thermal imaging camera footage (various models, Pulsar devices), depending on cave entrance orientation and observer distance. To assess population change, we compared the average counts per shelter in April 2024 and April 2026, calculating percent population change (Table 1).

**Table 1.** *Rousettus aegyptiacus* counts at surveyed shelters (April 2024 - April 2026).

| Shelter | Date |  |  |  |  |  | % Loss** |
| --- | --- | --- | --- | --- | --- | --- | --- |
|  | April 2024 | Dec 2025 | Jan 2026 | Feb 2026 | Mar 2026 | April 2026 |  |
| **Pafos** | 911-1037* | 330-356* | 3 | 5 |  | 5 | 99.5% |
| **Mammari** | 300 | 70 |  | 8 | 48 | 37 | 87.7% |
| **Pegeia** | 811-871* | 3 | 5 | 5 |  | 5 | 99.4% |
| **Pissouri** | 2518 | 138 | 184 | 355 | 415 | 411 | 83.7% |
| **Oreites** |  | 0 |  |  |  |  |  |
| **Androlykou** |  |  | 0 | 4 | 134 | 499 |  |
| **Petratis 1** |  |  | 0 |  |  |  |  |
| **Petratis 2** |  |  |  |  | 71 | 149 |  |
| **Mavrokolympos Dam** |  |  |  |  | 0*** |  |  |
| **Xylophagou\*\*\*\*** |  |  | 0 |  | 0 |  |  |
| **Episkopi \*\*\*\*** |  |  |  | 1 |  |  |  |
| **Total counted** | **4633** | **554** | **192** | **378** | **668** | **1106** |  |

\* Multiple counts/month; range: minimum-maximum.
\*\* Per site, % loss = 1 - (most recent monthly count / April 2024 count). If multiple site counts/month, use month's average.
\*\*\*Gated to exclude bats
\*\*\*\*SBA location

**Figure 1.**
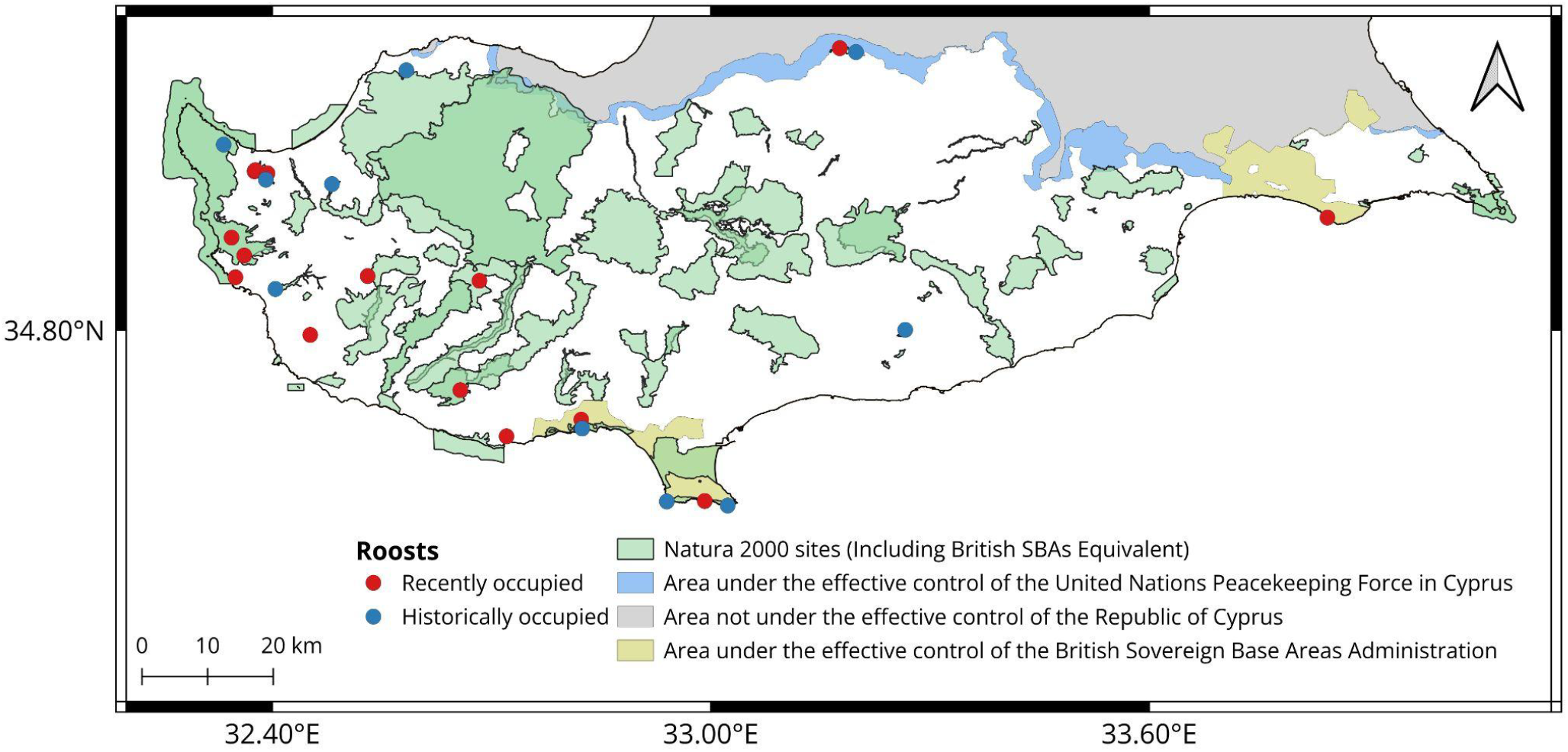
Map of Cyprus showing *Rousettus aegyptiacus* roosts. Table 1 roosts are “recently occupied.”

Results show that the Cypriot population of *R. aegyptiacus* experienced a rapid and severe population decline since April 2024. By December 2025, the *R. aegyptiacus* population in every cave assessed had declined by at least 64.8% and as much as 99.6%, and Oreites cave, occupied in 2022, was empty (Table 1). Many mummified dead bats were observed on three cave floors (Pegeia, Anavargos, and Xylophagou; Plate 1). Follow-up surveys in January through April 2026 recorded further evidence of decline (Table 1); no new dead bats were found. Total counts from all four original caves showed an overall decline of 90.1% over 24 months. This far exceeds qualifying conditions for Critically Endangered on the IUCN Red List (A2 Criteria: 80% decline over three generations or 15 years; Collen et al., 2016). The maximum count for each additional cave surveyed in 2026 was 499, 0, 149, 0, and one bat (Table 1).

The Cypriot *R. aegyptiacus* population has experienced previous population collapses. A Cypriot colonial government culling campaign beginning in the 1920s officially ended in 1993, and the bats received regulatory protection when Cyprus joined the European Union (Hadjisterkotis, 2006). A 2006 population of 10,000 bats declined to around 1,500 by 2010 (del Vaglio et al., 2011), reached 313 in 2017 (Lucan et al., 2026), then increased to 1000-1500 bats by 2018 (ECDR, 2018), 1050 bats in 2022 (Lucan et al., 2026), and 1000-5000 bats in 2024 (Reportnet3, 2026). The population is back to historically low levels (most recent count, 11 shelters: 1107 bats). Should the unknown cause of decline reoccur, there is likely not enough buffer in numbers to avoid extirpation.

The observed acute population decline is corroborated by evidence of a widespread mortality event. We observed many bat carcasses on three cave floors during December 2025 (Plate 1). Social media reported numerous dead and sick bats from February to April 2025 (Supplementary Material 1). An April 3, 2025 Cypriot Department of Environment announcement acknowledged high mortality (Ministry of Agriculture, 2025). Also, Pafos district farmers reported many *R. aegyptiacus* damaging fruit in 2024, but in 2025 both fruit damage and bat sightings had largely disappeared (A. Diez, personal communication, 2026). The evidence indicates that a major *R. aegyptiacus* mortality event occurred across the island.

**Plate 1.**
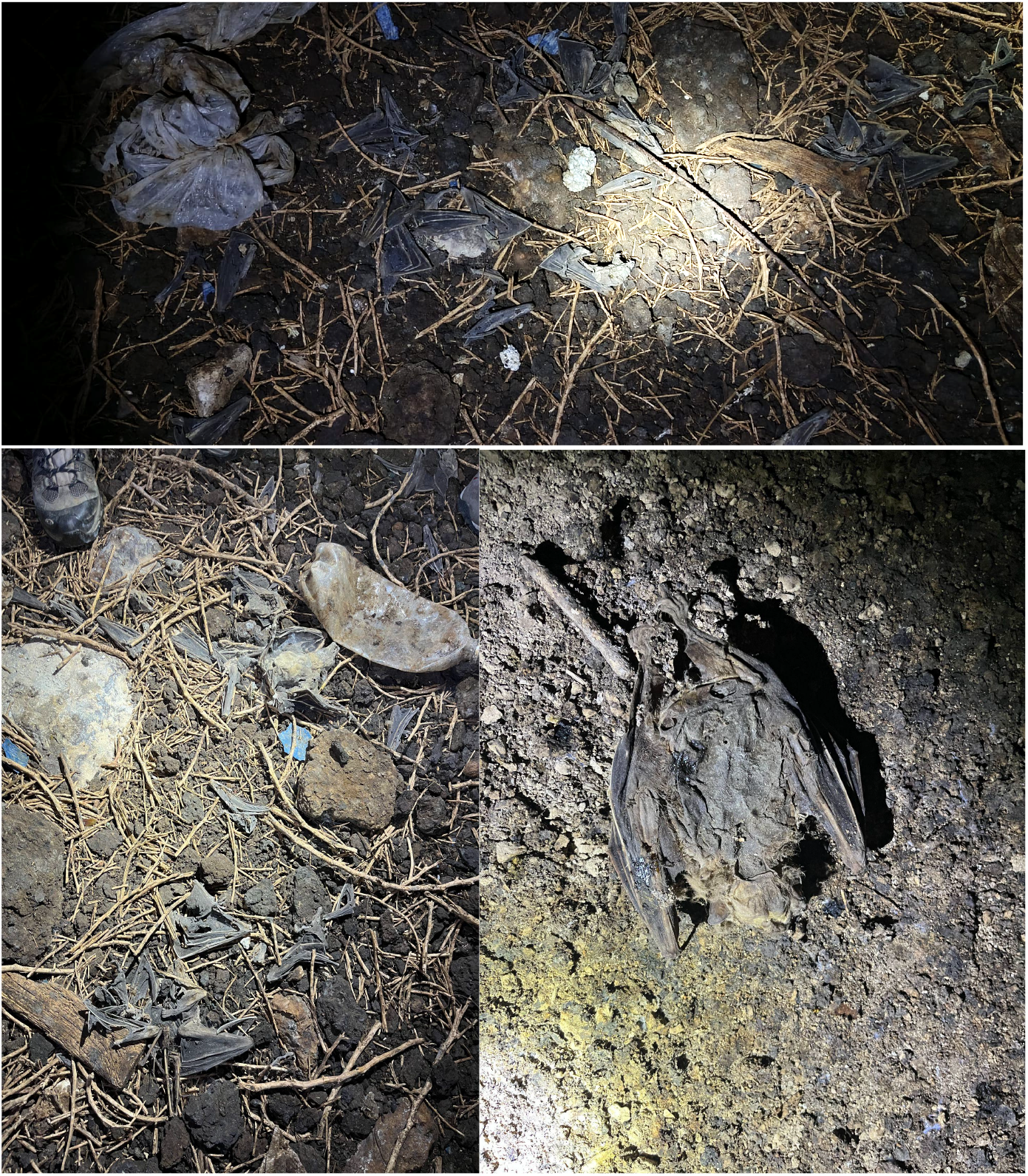
Dead *Rousettus aegyptiacus*, Cyprus; Kamarospilios (Pegeia) and Anavargos (Pafos) caves; December 4 and 5, 2025. Photos: Laura Fish, Dina Dechmann

Whereas some bats may have moved to alternative roost sites (not under effective control of the Republic of Cyprus), routine movements off the island are unlikely given this population’s genetic isolation (Hulva et al., 2012). Recent environmental stressors could change bat decision-making about long-distance flights, resulting in a regional metapopulation with movements on and off the island. However, in this case observed high mortality remains a serious conservation problem, functioning as a regional population sink (Dias, 1996).

Bat population decline can result from disease, habitat destruction and degradation, climate change, hunting, persecution, or combined factors (Russo & Dechmann, 2025). The cause of the current and past Cypriot *R. aegyptiacus* population declines is unknown. Threats considered potentially responsible include food shortage, pesticides, human conflict, and climate change (Russo & Cistrone, 2023; Lucan et al., 2026). In March 2025, carcass testing by a wildlife rescue agency identified *Staphylococcus aureus* (Supplementary Material 1), which has caused bat injury and death (Razik et al., 2024), but can be a sign of weakened immune function due to stress (Verbrugghe et al., 2012). The winter timing of mortalities could indicate food shortages. In Israel, low winter temperatures and food shortages contributed to annual cycles of increased morbidity in *R. aegyptiacus* (Weinberg et al., 2022). Cyprus has a regular seasonal climate, and the current decline does not appear to result from extreme weather or food shortages alone, as records show no extreme abnormalities (Supplementary Fig. 1). Without knowing the cause, urgent action is needed.

The magnitude, pace and ongoing nature of the decline warrant immediate coordinated action. Although the population has recovered from previous declines, a repeated collapse when numbers are still low might cause extirpation. We suggest establishing a joint task force of national authorities, local experts, and international advisors to coordinate emergency response actions. Ideally this task force should collaborate with experts in north Cyprus and the British base for island-wide protection. Actions should include rapid analysis of specimens to investigate causes of mortality, expedited permit processing for non-invasive monitoring and sample collection, regular emergence counts, national staff training in standardized monitoring and population estimation, and immediate protection of key roosts and foraging habitats until causes of the decline are understood and addressed.

## Author contributions

Study design, fieldwork: DD, AD, ZM, KM, MW; data analysis, writing: DD, KA, LW, AD, IM, ZM, KM, MTO, KP, MW, WF.

## Acknowledgements

This research received no specific grant from any funding agency, or commercial or not-for-profit sectors. Uschi Müller, Laura Fish, and University of Konstanz students helped with counts. Luis Víquez-R provided Figure 1. Kjell Wundram provided Supplementary Figure 1. We thank the Environment Department of the SBA Administration for support during fieldwork within SBA areas, facilitating access to survey sites, and issuing necessary permits under their legislation.

## Ethical standards

Data were collected with ethical permission from the Ministry of Agriculture, Agricultural Development and Environment, Cyprus, under permit numbers 02.15.001.003/04.05.002.005.006/02.15.007.003.002 (10/10/2022) and a SBA Administration Environment Department licence (Section 16, Protection and Management of Nature and Wildlife Ordinance, 2007), valid 12/1/2025-12/31/2026.

## Data availability

Data available within the article and supplementary materials.

