## Supplementary Materials for "Rapid population decline of Egyptian fruit bats on Cyprus"

**Supplementary Table 1. Roost distribution on the island of Cyprus** *Rousettus aegyptiacus* has been found in 25 roosts within the region under the jurisdiction of the Republic of Cyprus, the United Nations (UN) Buffer Zone and the Sovereign Base Areas (SBAs) of the United Kingdom between 2005-2025 (Hadjisterkotis 2006; ICOSTACY 2012; Nicolaou 2009, 2017, & 2024).

| Location | Number of roosts | Number of roosts that are protected and designated as Natura 2000 sites (or have equivalent legal status) |
|---|---|---|
| **Region under the jurisdiction of the Republic of Cyprus** | | |
| | 17 | 8* |
| **United Nations Buffer Zone** | | |
| | 2 | 2 |
| **British Sovereign Base Area** | | |
| | 6 | 6 |
| *Total* | 25 | 16 |

* Some important roosts are not covered by any formal protection (e.g. Pissouri, Pegeia, Polemi and Anavargos)

**Supplementary Material 1.** We list observations posted on iNaturalist (iNaturalist.org), social media posts, and personal communications documenting *Rousettus aegyptiacus* mortality in 2025: a) iNaturalist posts; b) social media group post summaries; c) personal communications with Klitos Papastylianou.

a) iNaturalist observations

Observed 17/02/2025 – Submitted 22/02/2025: https://www.inaturalist.org/observations/262737517

Observed 03/03/2025 – Submitted 03/03/2025: https://www.inaturalist.org/observations/263753803

Observed 08/03/2025 – Submitted 09/03/2025: https://www.inaturalist.org/observations/264660728

Observed 13/03/2025 – Submitted 19/03/2025: https://www.inaturalist.org/observations/265983757

Observed 20/03/2025 – Submitted 20/03/2025: https://www.inaturalist.org/observations/266137135

Observed 22/03/2025 – Submitted 22/03/2025: https://www.inaturalist.org/observations/266381937

Observed 17/04/2025 – Submitted 17/04/2025: https://www.inaturalist.org/observations/271010168

Observed 30/04/2025 – Submitted 03/05/2025: https://www.inaturalist.org/observations/278113383

Observed 30/04/2025 – Submitted 03/05/2025: https://www.inaturalist.org/observations/278150718

Observed 26/07/2025 – Submitted 31/07/2025: https://www.inaturalist.org/observations/302651067

b) Social media post summaries

- A March 16, 2025 post and comments in a private group included 11 people reporting finding between one and three dead Egyptian fruit bats on Cyprus. (Dechmann, Dina, personal communication, March 4, 2026).
- A January 2, 2025 post in a private group reported finding a dead Egyptian fruit bat on Cyprus (Dechmann, Dina, personal communication, March 4, 2026).
- A March 28, 2025 post in the public group “Cyprus Wildlife” at https://www.facebook.com/cypruswildlife/ [accessed March 4, 2026] reported receiving high numbers of sick and dead Egyptian fruit bats in February and March of 2025. The post also reported tests revealing *Staphylococcus aureus* in these bats. (Dechmann, Dina, personal communication, March 4, 2026).

c) Personal communications

Photos courtesy of Klitos Papastylianou taken on February 20, 2025. According to the photographer, two dead fruitbats were observed close to a roost of the species, in Avakas Gorge on Akamas Peninsula. Several more dead bats were found in the gorge and the broader area in the same period.

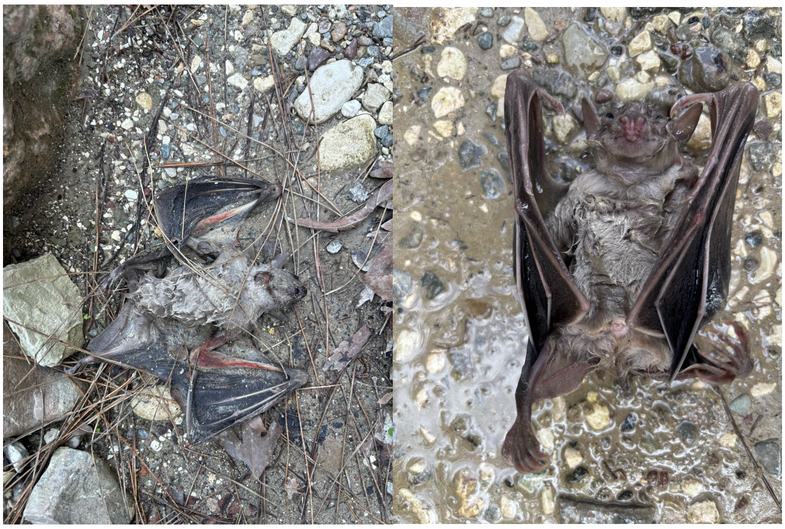

**Supplementary Figure 1.** Seasonal patterns on Cyprus from 1999–2026 showing land surface air temperature at 2 m in °C (top; Muñoz Sabater 2019), monthly precipitation totals in millimetres (middle; CHIRPS3 2025), and Normalized Difference Vegetation Index (NDVI) as an indicator of vegetation greenness and primary productivity (bottom; Didan 2021). Solid lines show the island-wide median, and shaded bands represent the spatial interquartile range across Cyprus.

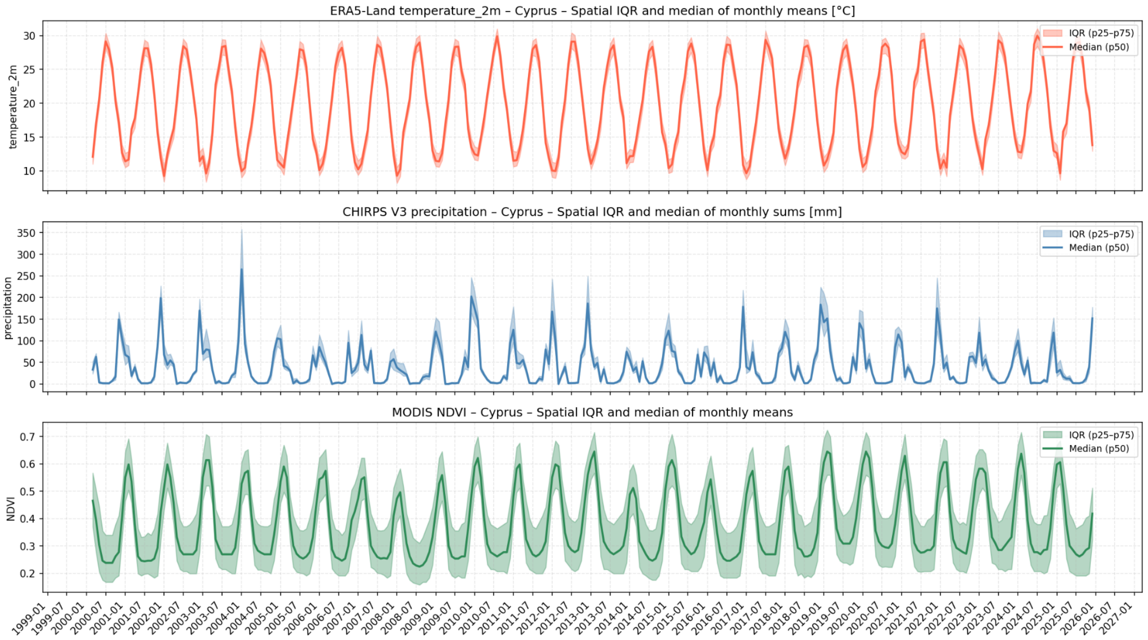
